# Framing the potential of public frameshift peptides as immunotherapy targets in colon cancer

**DOI:** 10.1101/2020.03.26.010231

**Authors:** Ide T. Spaanderman, Fleur S. Peters, Aldo Jongejan, Egbert J.W. Redeker, Cornelis J.A. Punt, Adriaan D. Bins

## Abstract

Approximately 15% of Colon Cancers are Microsatellite Instable (MSI). Frameshift Peptides (FPs) formed in MSI Colon Cancer are potential targets for immunotherapeutic strategies. Here we comprehensively characterize the mutational landscape of 71 MSI Colon Cancer patients from the cancer genome atlas (TCGA-COAD). We confirm that the mutations in MSI Colon Cancers are frequently frameshift deletions (23% in MSI; 1% in microsatellite stable), We find that these mutations cluster at specific locations in the genome which are mutated in up to 41% of the patients. We filter these for an adequate variant allele frequency, a sufficient mean mRNA level and the formation of a Super Neo Open Reading Frame (SNORF). Finally, we check the influence of Nonsense Mediated Decay by comparing RNA and DNA sequencing results. Thereby we identify a set of 20 NMD-escaping Public FPs (PFPs) that cover over 90% of MSI Colon Cancer patients and 3 out of 4 Lynch patients in the TCGA-COAD. This underlines the potential for PFP directed immunotherapy, both in a therapeutic and a prophylactic setting.

## Introduction

Protein products derived from mutated regions of DNA are a source of neo-antigens that can drive immune activation against cancer and convey susceptibility to immunotherapy^1–4^. Specifically, frameshift insertions and deletions (INDELS) are able to drive the anti-tumor immune response^5^. INDELS in coding DNA result in frameshifted RNA containing neo open reading frames (NORFS) that, once translated, give rise to completely non-self, out-of-frame protein products. These out-of-frame proteins can be degraded into peptides and presented in the context of multiple MHC alleles. Recognition of these MHC-peptide complexes by CD4+ or CD8+ T-cells initializes an anti-tumor immune response and leads to an influx of T cells into the tumor microenvironment^1,6,7^, Longer NORFS (so called super NORFS: SNORFS), have a higher chance to be recognized as a neo-antigens^8^. Importantly, NORFS are subject to degradation on a RNA level by the Nonsense Mediated Decay (NMD) pathway^9^. In line with this, the presence of NMD escaping SNORFS has been shown to be a strong predictor of immunotherapy response and survival in multiple cancer types^8^.

Compared to other mutation signatures, INDELS are overrepresented in the exome of microsatellite instable (MSI) tumors^5,10^. Microsatellites are regions in the DNA with a repeated nucleotide motif. These regions are prone to polymerase slippage and subsequently incorrect re-annealing during DNA synthesis. This leads to single stand insertions and deletions in the nascent DNA strand^11^. In healthy cells, the mismatch repair (MMR) system recognizes and repairs these damages. However, MSI tumors have a deficient MMR system (dMMR) due to germline and/or acquired somatic mutations in one or multiple MMR genes (*MLH1*, *MSH2*, *MSH6*, *PMS2*, *EPCAM*) or by hyper methylation of the *MSH1* promotor^12,13^. Approximately 15% of colon cancers are considered MSI^14^. These dMMR colon cancers have high numbers of mutations, specifically INDELS in microsatellites.

Here we comprehensively asses frameshift mutations in colon cancer samples sequenced within the genome atlas (TCGA-COAD)^15^. We focus on the TGCA-COAD series as this contains a large subset of MSI tumors. Using a novel in silico approach (**figure 1**), we establish that colon cancer frameshift mutations cluster to specific loci in the colon cancer exome. Based on DNA and RNA sequencing data we investigate whether these mutations lead to SNORFS and subsequently whether these SNORFS are affected by NMD. Finally, we identify a set of frequently occurring frameshift peptides (FPs), that we name public frameshift peptides (PFPs). These PFPs are potential targets for immunotherapy strategies in an adjuvant setting, as they obviate the need for time-consuming personalization of vaccines. Moreover, PFPs can also be found in all four (Lynch) patients with a germline MMR mutation in the TCGA-COAD. This highlights the possibility for PFP specific vaccination in a prophylactic setting as well.

**Figure 1.**
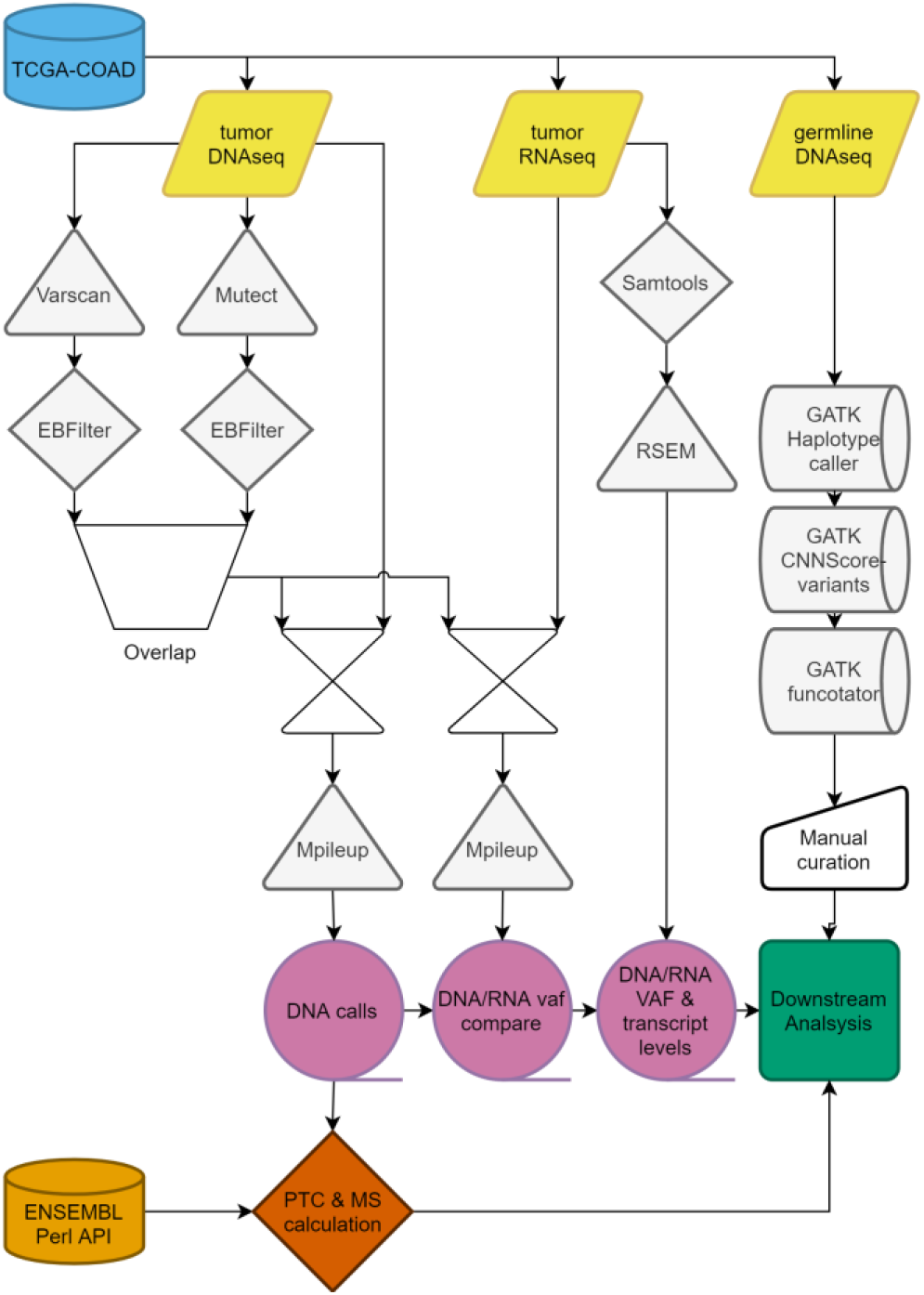
Schematic overview of in silico pipeline. TCGA-COAD: the cancer genome atlas colorectal adenocarcinoma database; Varscan: somatic variant calling; Mutect: somatic variant calling; EBFilter: Bayesian filtering for somatic variants; Mpileup: location specific read calling; RSEM: transcription quantification; VAF: variant allele frequency; ENSEMBL Perl API: Programmable interface for Ensembl genome browser; GATK: Genome analysis toolkit; Haplotypecaller: genetic variant calling; CNNScoreVariants: germline variant filtering; Functotator: Clinvar based annotator of genetic variants; PTC: premature termination codon; MS: microsatellite

## Results

### Frameshift mutations cluster to specific genomic loci in MSI colon cancer

In line with previous reports in literature 71 out of 461 (15%) of TCGA-COAD patients are MSI^14^. As expected, MSI colon cancer patients have a higher mutational burden than microsatellite stable (MSS) colon cancer patients (MSI: median = 2974, n = 71; MSS: median = 218, n = 272; p < 0.0001) (**figure 2a**) and a larger percentage of their mutations consists of deletions or insertions (MSI: SNP = 72%, INS = 5%, DEL = 23%’; MSS: SNP = 97%, INS 1%, DEL = 1%) (**figure 2b**). Almost all of these INDELS are frameshift mutations (INS: Frameshift 99%; DEL: Frameshift 98%) (**figure 2c**). Contrary to single nucleotide polymorphisms (SNPs), these INDELS are preferentially located in coding microsatellites (cMS), pinpointing the role of the dysfunctional MMR system in the mutation signature of MSI colon cancer (INS: median = 7; DEL: median = 7; SNP: median = 1, p < 0.0001) (**figure 2d**). As expected, both SNP and INDEL mutations cluster to specific loci in the MSI colon cancer exome. Similar to the well documented BRAF SNP at chr7:140753336^16^, it seems probable that the most prevalent INDEL mutations also play a role in tumorigenesis, (**figure 3a**). 13 of the 14 most prevalent mutations are frameshift mutations located in a cMS. Strikingly, the most frequently occurring frameshift mutation is present in 46% of the patients. 123 frameshift mutations occur in more than 10% of the patients (**figure 3b**). Notably 82.5% of the deletions that occur in more than 10% of patients are listed in the candidate cancer gene atlas and 42.5% are listed as potential drivers in colon cancer^17^ (**supplementary table 1**). Interestingly, when comparing the tendency to cluster among DEL, INS and SNPs mutations, clustering is only clearly present for DEL mutations (median percentage of patients with a similar specific mutation; DEL = 14.08%, INS = 4.23%, SNP = 4.23%, p < 0.0001) (**figure3c**).

**Figure 2.**
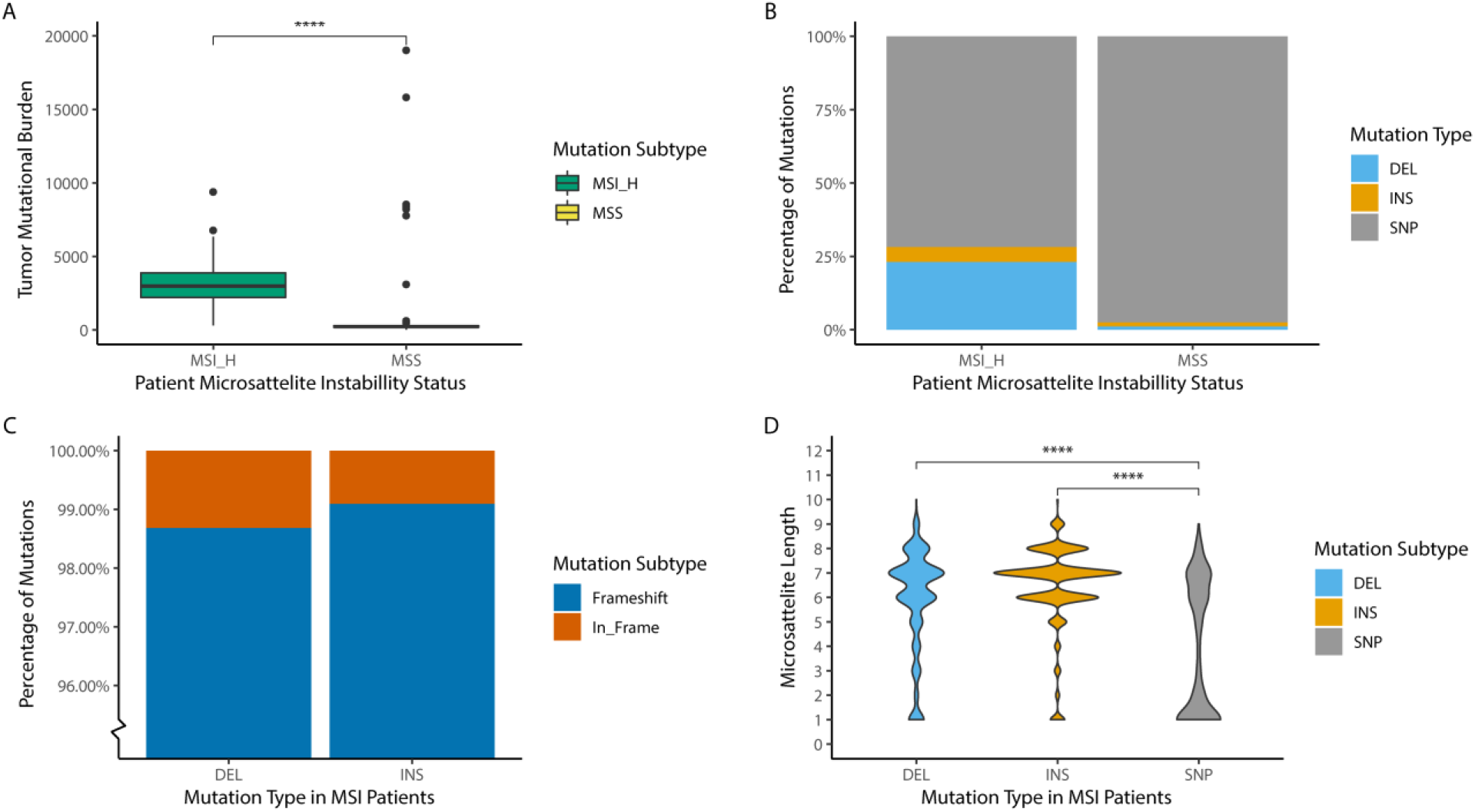
Mutational profile for MSS and MSI in TCGA-COAD. **A)** Difference in tumor mutational burden as number of mutations per patient between MSI (median = 2974, n = 71) and MSS patients (median = 218, n = 272, p < 0.0001). **B)** Mutation types as percentage of mutations between MSI (DEL = 23.15%, INS = 5.06%, SNP = 71.19%, n = 71) and MSS patients (DEL = 1.21%, INS = 1.34%, SNP = 97.4%, n = 218) C) Percentage of frameshift mutations for deletion (Frameshift = 98.69%, In_Frame = 1.31%, n = 53851) and insertions (Frameshift = 99.07%, In_Frame = 0.93%, n = 13172) in MSI patients (n = 71). D) Difference in microsatellite length in coding mononucleotide repeats on location of mutation in the genome between mutation subtypes, DEL (median = 7, n = 4960), INS (median = 7, n = 2384), SNP (median = 3, n = 1859, p < 0.0001) in MSI patients (n = 71). Microsatellite length of 3 mononucleotide repeats or less cannot be considered true microsatellites.

**Figure 3.**
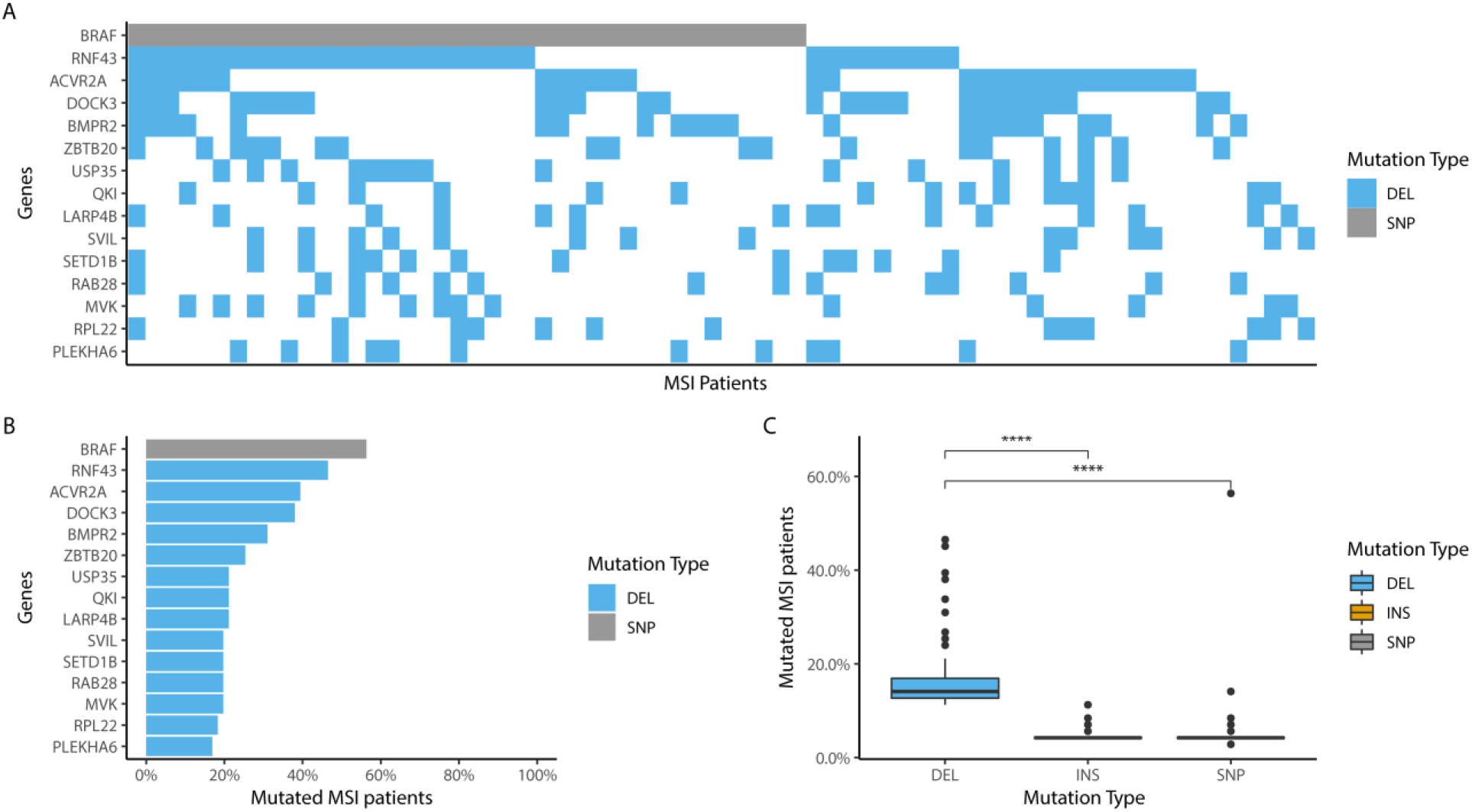
Mutation clustering in MSI TCGA-COAD. **A)** Waterfall graph depicting the 15 most frequently occurring mutation hotspotsby Gene Identifier in MSI patients. Light grey mutations are SNPs. Light blue mutations are DELs. Other mutation types do not occur within the 15 most frequently occurring mutation hotspot. **B)** Percentage of most frequently occurring mutation loci. **C)** Difference in mutation frequency as percentage of patients with mutation on locus between mutation types for the top 100 most frequently occurring mutations, DEL (median = 14.08%, n = 100), INS (median = 4.23%, n = 100), SNP (median = 4.23%, n= 100, p < 0.0001) in MSI patients (n = 71).

### Nonsense mediated decay escape

In light of specific immunotherapy approaches it is important to further investigate the identified public frameshift mutations. As previously mentioned, NORFS derived from INDELS are potentially subject to NMD. NMD is triggered by the formation of a premature termination codon (PTC), that is prone to occur in out-of-frame NORFS^9^. If such a PTC is located at least 50 base pairs before the last exon-exon junction, the mRNA is considered sensitive to degradation by NMD. If the PTC is located behind this point the transcript is considered NMD resistant^18^. In the MSI cancers of the TGCA-COAD this concept holds true, as the mean z-score for mutated transcripts with a NMD sensitive PTC is significantly lower than that of a mutated transcripts with a NMD resistant PTC (median SENS = −0.4655; RES = −0.0657; p < 0.0001) (**figure 4a**). Nonetheless the NMD effect is limited in an absolute sense and clearly heterogeneous as a large portion of potential sensitive mutations seem to escape NMD based on the Z-score of mRNA transcript levels. However, the fact that a single specific mutation can be mapped to multiple transcripts (isoforms), might limit the sensitivity for detection of transcript specific NMD signals.

**Figure 4.**
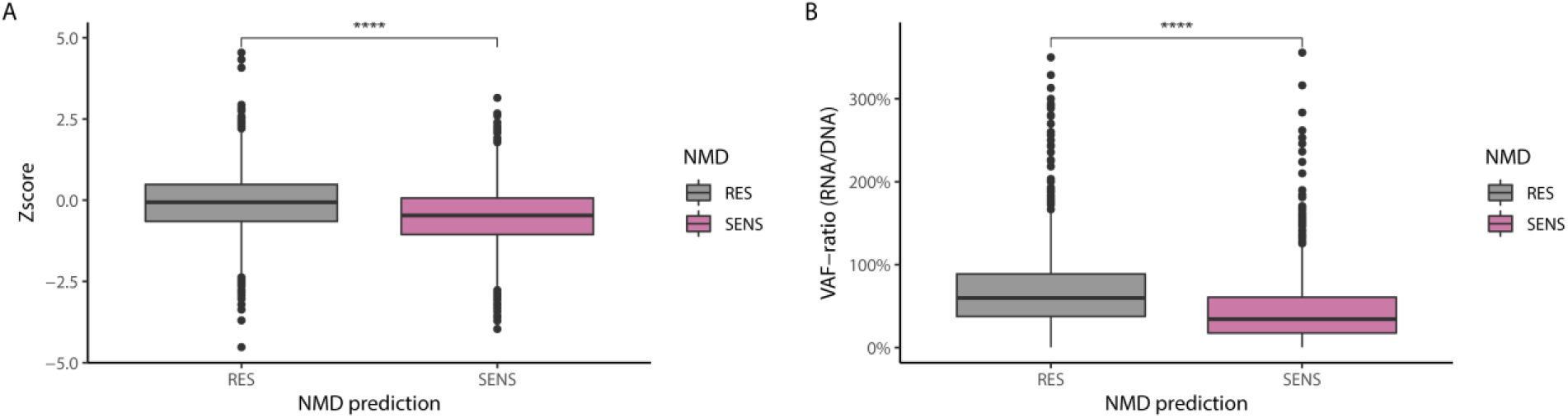
Nonsense Mediated Decay Escape in MSI TCGA-COAD. Approximation of Nonsense Mediated Decay(NMD) on mutated transcripts either predicted as NMD sensitive (SENS) or NMD resistant (RES), based on the location of the premature termination codon (PTC): RES PTC is located 50bp or less before the last exon-exon junction or in last exon **A)** NMD defined as a z-score of transcript levels for RES (median = −0.0657, n = 1357) and SENS (median = −0.4655, n = 1024, p < 0.0001). **B)** NMD defined as VAF-ratio: the allele frequency (VAF) as percentage RNA VAF of DNA VAF for RES (median = 59.9%, n = 936) and SENS (median = 34.3%, n = 904, p < 0.0001).

To negate this problem, we checked the Variant Allele Frequency (VAF) for each mutated locus in the DNA and RNA. The VAF is the percentage of mutated reads in all reads on that specific locus. We calculated the difference between the DNA VAF and RNA VAF: the VAF-ratio. This transcript independent method is a much more sensitive and accurate measurement of NMD (**Supplementary figure 1**). On average, for NMD sensitive mutations the variant allele frequency (VAF) in RNA sequencing data is decreased by 65.7% compared to the DNA VAF (median VAF-ratio 34.3%) n = 904), which is significantly more than the 40.1% decrease in NMD resistant mutations (median VAF-ratio 59.9%, n = 936; p < 0.0001) (**figure 4b**). This clearly confirms that NMD is an important factor in the formation of neo-antigens from NORFS in MSI colon cancer. Nonetheless roughly one third of NMD sensitive NORFS still escape NMD, with a VAF-ratio of 50% or more. This presents a clear opportunity for PFP formation.

### NMD escape by public frameshift peptides

Besides the potential NMD impact of PTCs, PTCs also can limit NORF length to a size that prohibits presentation in the context of MHC. In the NORFS identified by us in MSI cancers of the TGCA-COAD, a PTC occurs on average after 14 amino acids (**Supplementary figure 2**). Therefore, we selected potential PFP by discarding NORFS with a length below 10 amino acids. Hereby, we identified 90 SNORFS occurring in more than 5% of MSI patients. Assuming that a set of 20 PFPs is feasible to incorporate in a multivalent vaccine, we discarded the 70 least frequently occurring SNORFS (**table 1**). The remaining 20 frequently occurring SNORFS have a mean NORF length of 28 amino acids, a mean VAF-ratio of 67.93% and a mean mRNA level of 2.30 (Log2 TPM) for the most abundantly available corresponding transcript (isoform). Most importantly, at least one of these 20 SNORFS is present in the exome of 93% of MSI samples in the TGCA-COAD. A limited set of 10 SNOFRS, as has been used in a personalized RNA vaccination setting^19^, would still cover 74.6% of these patients (**Supplementary figure 3**). This highlights the potential for non-personalized immunotherapy that targets PFP in MSI colon cancer patients. Notably, 25% of these 20 SNORFS occur are listed as potential drivers in colon cancer^17^.

**Table 1.**
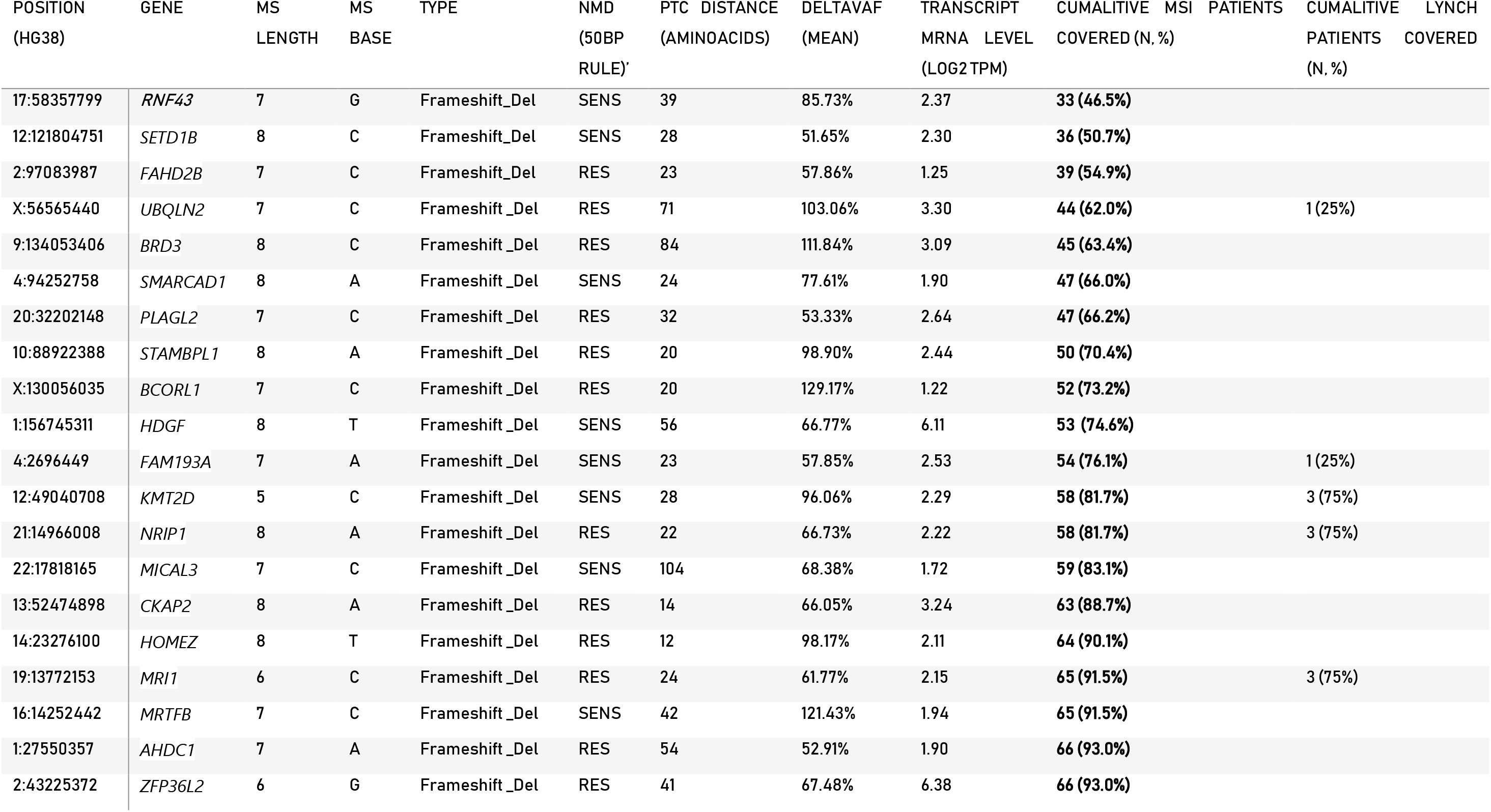
Public Frameshift Proteins (PFP). Properties of the 20 most frequently occurring mutations with SNORF of at least 10 Amino Acids and DeltaVAF of 50%. MS: microsatellite. Bold: cumulative numbers of MSI patient covered by mutation (n = 71). Germline patients n = 4. NMD: predicted NDM status based on PTC location in transcript. DeltaVAF: mean RNA-VAF percentage of DNA-VAF for all mutated patients. Expression: log2(TPM + 1) for most abundant corresponding transcript..

### PFP set covers germline mismatch repair deficient tumors in TCGA-COAD

Lynch patients carry a heterozygous germline mutation in an MMR gene, resulting in a 50-80% life-time chance to develop colon cancer^20^. MSI colon cancer tumorigenesis in Lynch patients may be different than that of sporadic MSI colon cancer, leading to expression of different PFPs. At the same time, prophylactic immunotherapy targeting PFP may be beneficial for these patients. Therefore we evaluated whether the PFP set may potentially be applicable to patients with a germline MMR mutation. Using normal blood DNA sequencing data, we identified 4 MSI TCGA-COAD patients (5.6%) with a pathogenic germline dMMR mutation. This percentage is in line with earlier reports of germline dMMR mutations in literature^20^. In overlap with sporadic MSI colon cancers, three out of four of these germline mutated dMMR tumors encodes at least 1 PFP from the 20-valent PFP-set. Colon Cancer samples with a germline MMR mutation from two patients only have 1 mutation in one of the identified hotspots, but 1 germline mutated MSI sample even has 4 hotspot mutations (**table 1**). This supports the potential of the PFP-set in a prophylactic vaccination setting for Lynch patients.

## Discussion

Based on DNA and RNA sequencing data from MSI colon cancer patients in the TGCA we identified a subset of frameshift mutations that cluster to distinct cMS in the genome, produce SNORFS that evade NMD, and have sufficient length for presentation in MHC context. Based on this, we established a PFP-set of 20 frameshift peptides that cover 93% of individual MSI patients. Such a set of PFPs has high potential for non-personalized cancer-specific immunotherapy strategies directed against MSI colon cancer, especially in the adjuvant setting. In addition, this set of PFPs also covers three out of four Lynch patients in the TGCA-COAD, which supports potential application in a prophylactic vaccination setting for patients with Lynch syndrome. Cleary, as the sporadic MSI colon cancer mutation signature may differ from that of Lynch patients, target discovery for Lynch patients requires a larger patient cohort than the 4 patients identified in the TCGA-COAD.

We did not predict MHC binding for the selected PFPs. Currently MHC class I binding predictions tools only achieve approximately 40% sensitivity^21^. Although new methods have recently been published^22^, especially the prediction of MHC class II binding remains very challenging, whilst MHC class II presented neo-antigens seem to drive anti-tumor immunity^22,23^. Furthermore, the above average length of the PFPs and the fact that they are non-self on any position makes adequate presentation within a naturally polymorphic MHC context more likely. Also, the concept of non-personalized immunotherapy would be compromised by MHC matching. In this context it is noteworthy that the loss of MHC-I on colon cancer cells could potentially limit the effectiveness of immunotherapy that target PFPs in the same way it can limit the effectiveness of checkpoint inhibition.^24^

Finally, since this work is based on in silico analysis of sequencing data we could not determine the occurrence of PFPs on a protein level. Nonetheless, based on our extensive filtering for NMD signals in RNA sequencing data we deem it likely that a majority of the selected SNORFS will be translated and lead to presentable PFPs. Alsco it has been shown that a first so-called pioneering round of translation, can already be adequate for neo-antigen formation^25^. In that regard our higher VAF threshold and strict NMD evasion filtering might even hinder identifying relevant PFPs. Furthermore the availability of RNA sequencing data also negated the need for novel NMD prediction algorithms^26^. In order to assess the FPF presentation on a protein level we are currently collecting patients samples in the ATAPEMBRO trial^27^, which we aim to screen for specific PFP T-cell immunoreactivity by IFN-gamma Elispot.

In conclusion we identified a set of 20 DEL mutation hotspots in coding DNA microsatellites that give rise to adequately transcribed, NMD-escaping SNORFS and subsequently can produce PFPs. Likely, these SNORFs are involved in colon cancer tumorigenesis. The 20 most common PFPs cover 93% of MSI patients and 3 out of 4 patients that harbor a pathogenic germline MMR mutation in the TCGA-COAD. Therefore, these PFPs are potential targets for immunotherapy strategies in MSI colon cancers.

## Methods

### Data acquisition, storage and computational analysis

All DNA and RNA sequencing data and corresponding patient characteristics, microsatellite panel and *MLH1* promotor methylation data was collected from the Cancer Genome Atlas Colon Adenocarcinoma Database (TCGA-COAD)^15^. Access to private data was granted by the National Cancer Institute Data Access Committee (NCIDAC) from the U.S. National Institute of Health trough the database of Genotypes and Phenotypes (dbGaP) portal. According to dbGaP guidelines private data was stored on the pre-approved internal Amsterdam UMC data-infrastructure. Computations on large public TCGA datasets were performed on the SurfSara high performance computing cloud (grant provided by the Dutch government).

### Somatic mutation calling from DNA sequencing files

Somatic mutations were called on aligned reads by Mutect^28^ and Varscan^29^ separately with default quality control parameters. Those calls were independently filtered by empirical Bayesian mutation filtering using EBFilter^30^ to reduce the false positive rate. Finally, the overlapping filtered mutations between these two methods for each individual patient were selected as true positive somatic mutations. VAF was calculated as percentage of total reads that contains mutation. A VAF cutoff of 20% was used.

### Nonsense mediated decay escape by RNA and DNA mutated reads comparison

Counts for mutated reads were collected by performing Mpileup^31^ on matched RNA and DNA tumor aligned sequencing reads for each of the specific genomic position of the somatic mutations selected prior. Mutations with less than 10 mutated reads in the DNA were discarded, although this number was very limited due to the stringent selection of mutations. No lower limit was placed on the RNA read counts since this would hinder the detection of complete nonsense mediated decay (NMD). NMD is measured by the difference in percentage between the fraction of mutated reads in the DNA compared to the RNA (or ΔVAF), were the fraction of mutated reads in the DNA is set as 100%.

### Transcript mRNA levels and Zscore

Accurate transcript quantification from RNA sequencing data is calculated by RSEM^32^ and mapped to Ensembl^33^ genome browser transcripts from reference genome GRCh38.p13. Absolute mRNA transcript levels per million (TPM) are 2log transformed. The z-score for mRNA levels of each mutated transcript in a specific patient is calculated based on the following equation for transcript with a mean mRNA level of more than 1.0 in all patients, removing transcripts that are not expressed in colon cancer tissue.

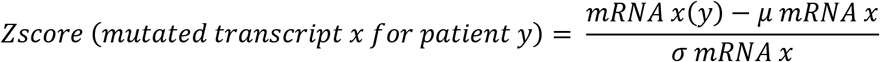

### Microsatellite and premature termination codon calculation

A proprietary Perl script run with GNU parallel utilizes the Ensembl Perl API to collect transcript sequences based on the position, allele, type and length of inputted true positive somatic mutations. Using this sequence, it calculates the number of mono-, di- or multi-nucleotide repeats on the mutation position. Subsequently it constructs the new reading frame after mutation and checks the RNA sequence until the occurrence of a premature termination codon. Finally, it records the position of this PTC in the transcript and relative to the mutational site.

### Germline mutation calling from normal tissue DNA sequencing files

Aligned DNA read files were spliced to incorporated only genomic regions of the mismatch repair genes *MLH1*, *MSH2*, *MSH6*, *PMS2* and *EPCAM*. Germline mutation calling on spliced aligned reads was performed using the GATK germline short variant discovery pipeline and using the GATK^34^ best practices. The workflow subsequently utilizes Haplotypecaller, CNNScoreVariants and FilterVariantTrances. Mutation calls with QD < 10 and read count < 10 were discarded. Mutational calls were functionally annotated employing Funcotator^34^. Variants of unknown significance were manually scored by a laboratory specialist clinical geneticist on a 1-5 scale according to the American College of Medical Genetics and Genomics (ACMG) guidelines^35^, based on mutation characteristics, amino-acid change and taking patient clinical data, somatic mutations and MLH1 promotor methylation data into account.

### Downstream analysis

Statistical analysis were performed using R 3.4.4. in Rstudio 1.2.1335 (Rstudio, INC) with the tidyverse package enabled. Difference between means are calculated by unpaired t test for comparison between 2 groups and ANOVA for more than 2 groups. Graphs were generated using ggplot2.

## Acknowledgments

This work is made possible by the Dutch Cancer Society (KWF) Grant 21923. We would like to thank Prof. dr. L. Vermeulen for proofreading the manuscript.

## Author Contributions

I.T.S and A.D.B. conceived the main hypothesis. I.T.S, F.P and A.D.B designed the experiments, analyzed the data and wrote the paper. I.T.S. developed the code and performed the experiments. I.T.S. and A.J. set up the in silico testing environment. I.T.S and B.R identified samples with a germline MMR mutation. A.D.B and C.J.A.P. supervised the project. All authors read and commented on the manuscript.

## Competing Interests

The authors declare no competing interests.

## Code Availability

The proprietary Perl computer code used to calculate PTC position in coding microsatellites is available at https://github.com/ITSpaanderman/PTC_predictor, including run instructions and test samples. GATK pipeline information is available at https://gatk.broadinstitute.org/hc/en-us. RSEM computer code is made available by its author at https://github.com/deweylab/RSEM.

## Supplementary Figures

**Supplementary figure 1.**
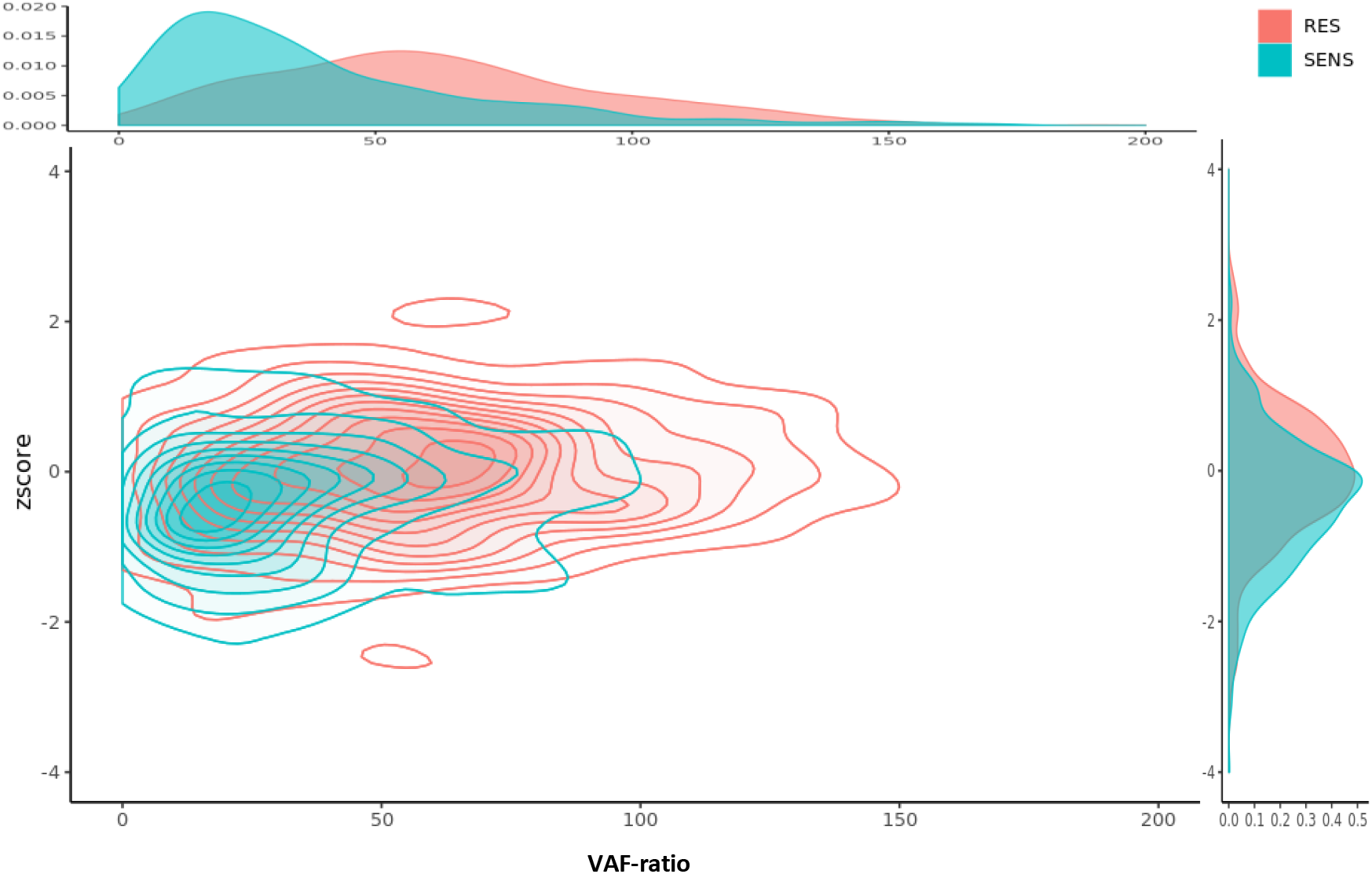
Difference in NMD measurement. Density map showing difference between Z-score and VAF-ratio approach for the measurement of Nonsense Mediated Decay (NMD) in NMD RES and SENS mutations. Separation of NMD RES and SENS is clearer with VAF-ratio.

**Supplementary figure 2.**
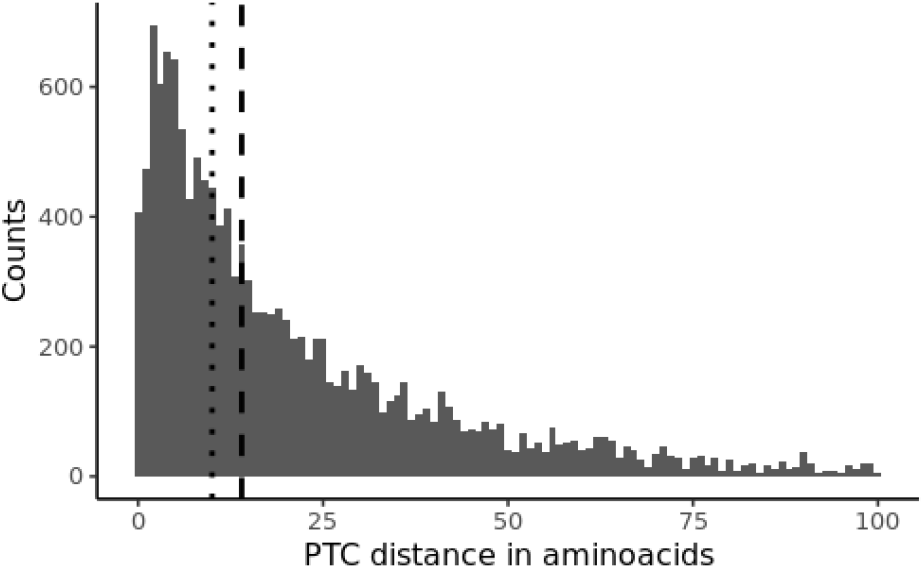
Premature Termination Codon (PTC) distance. PTC distance for mutation in MSI_H. Median is shown as dotted line (median = 14 Amino Acids). Dashed line shows 20 Amino Acids.

**Supplementary figure 3.**
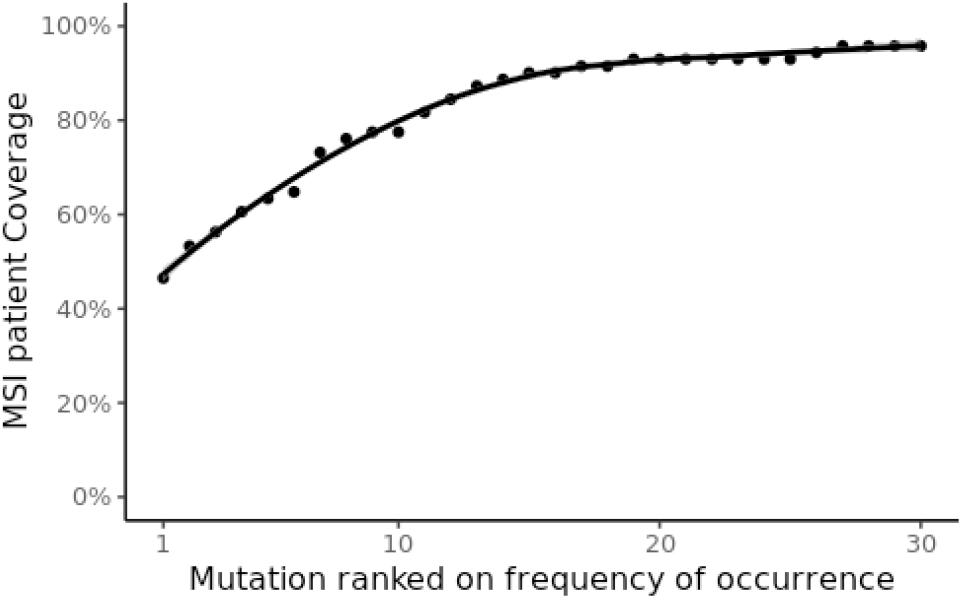
Patient coverage for top 30 frameshift mutations. Percentage of patients covered by n most frequently occurring frameshift mutations with adequate expression, a SNORF and NMD-escape.

## Supplementary Tables

**Supplementary Table 1.**
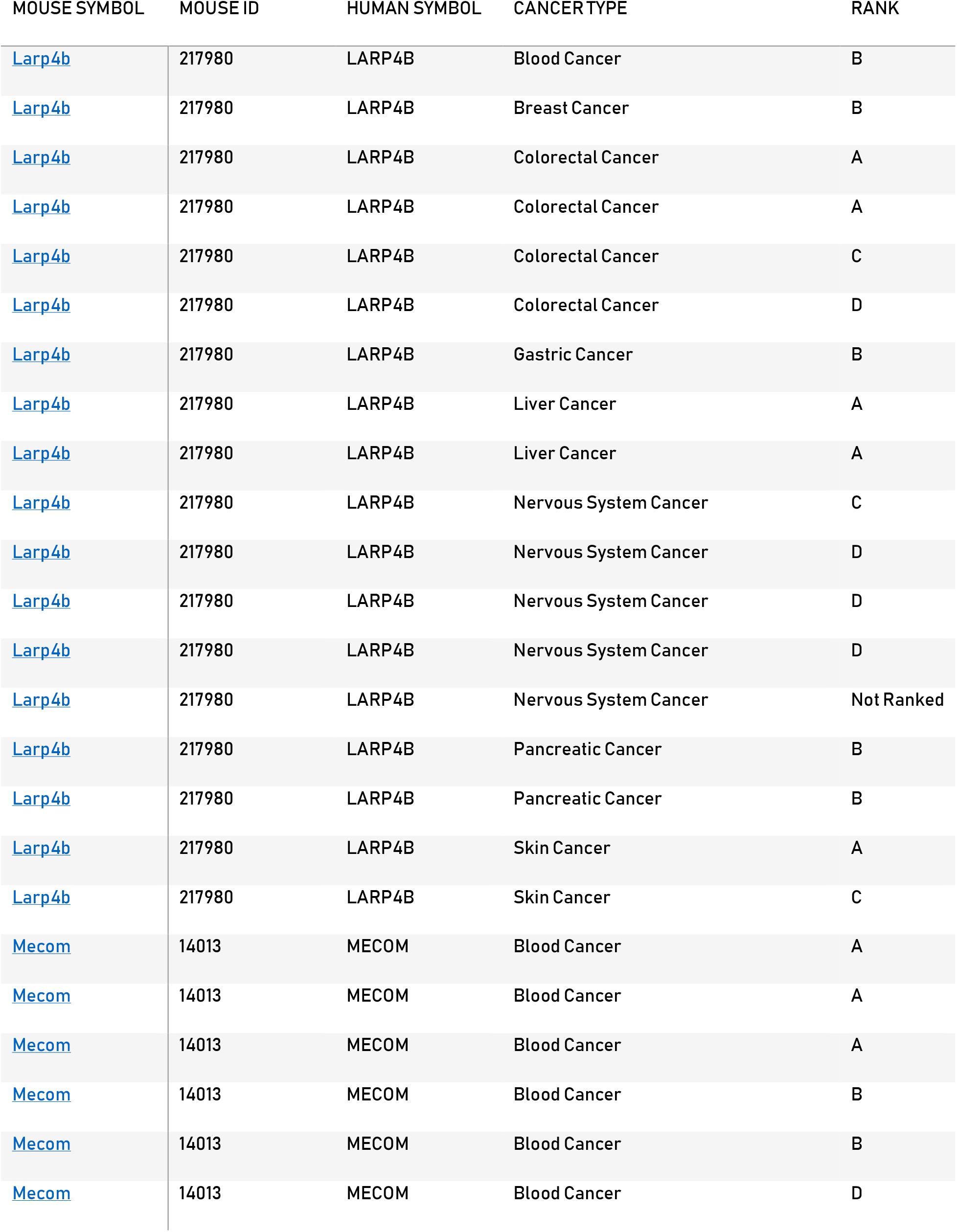
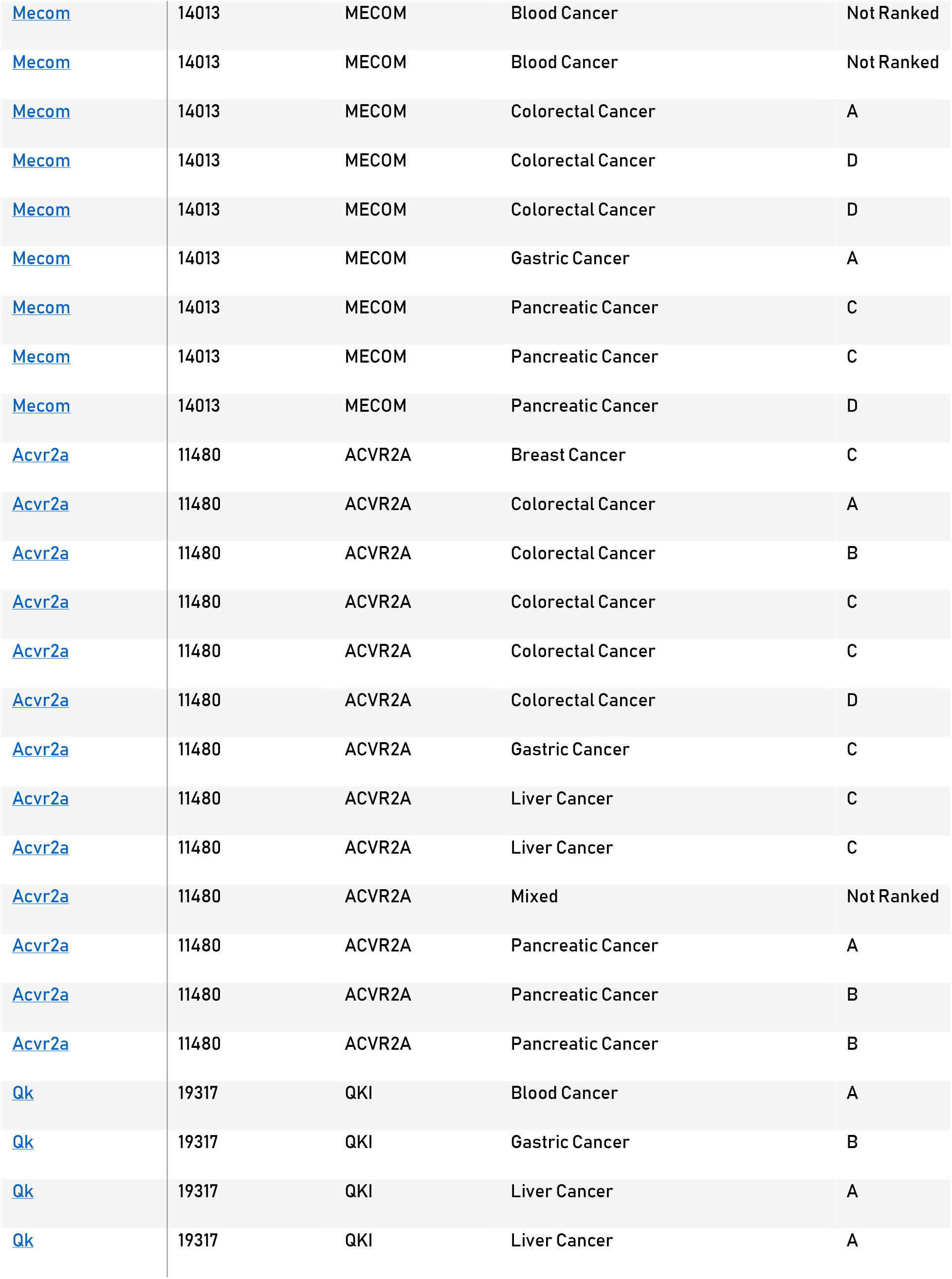
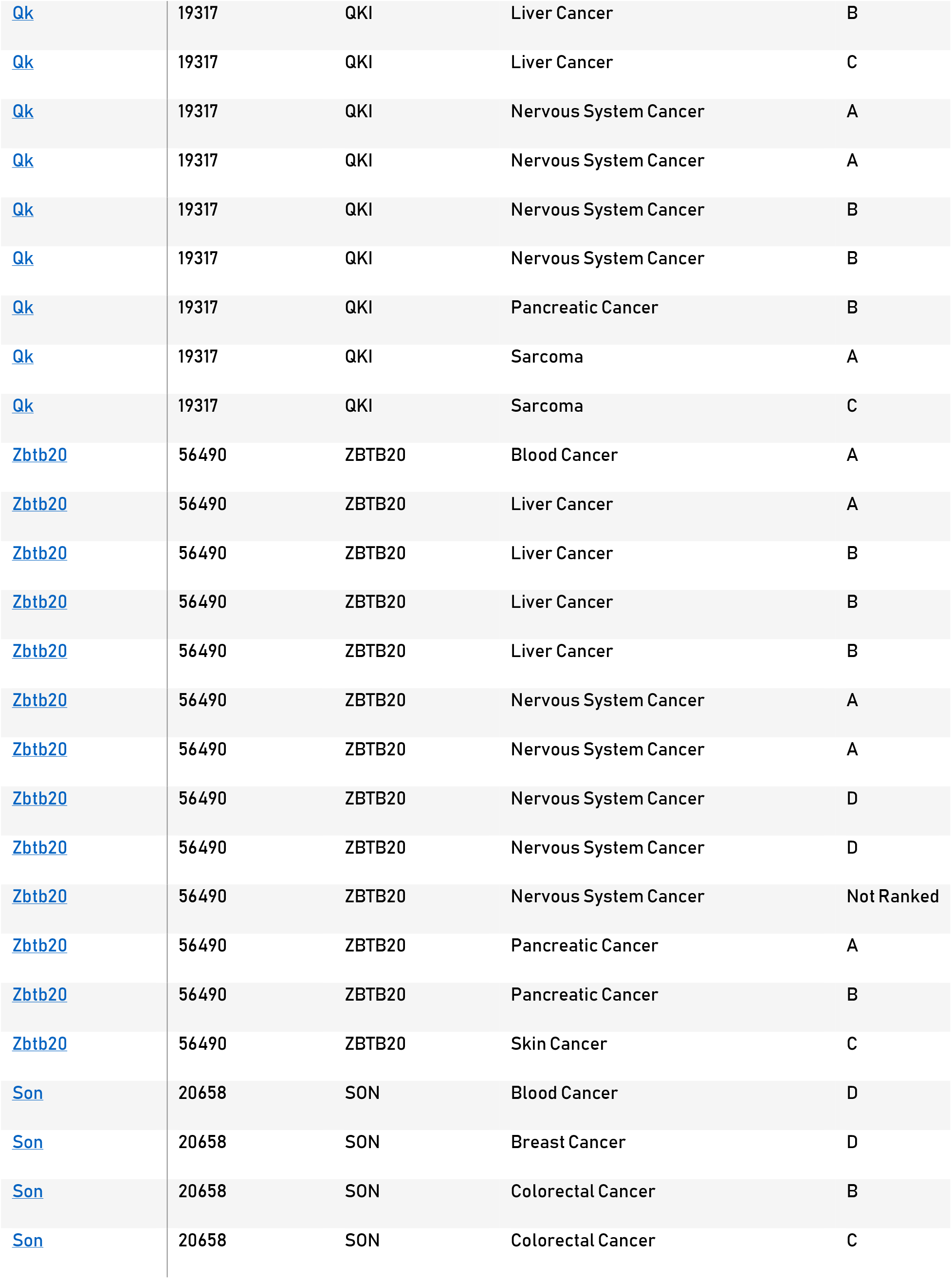
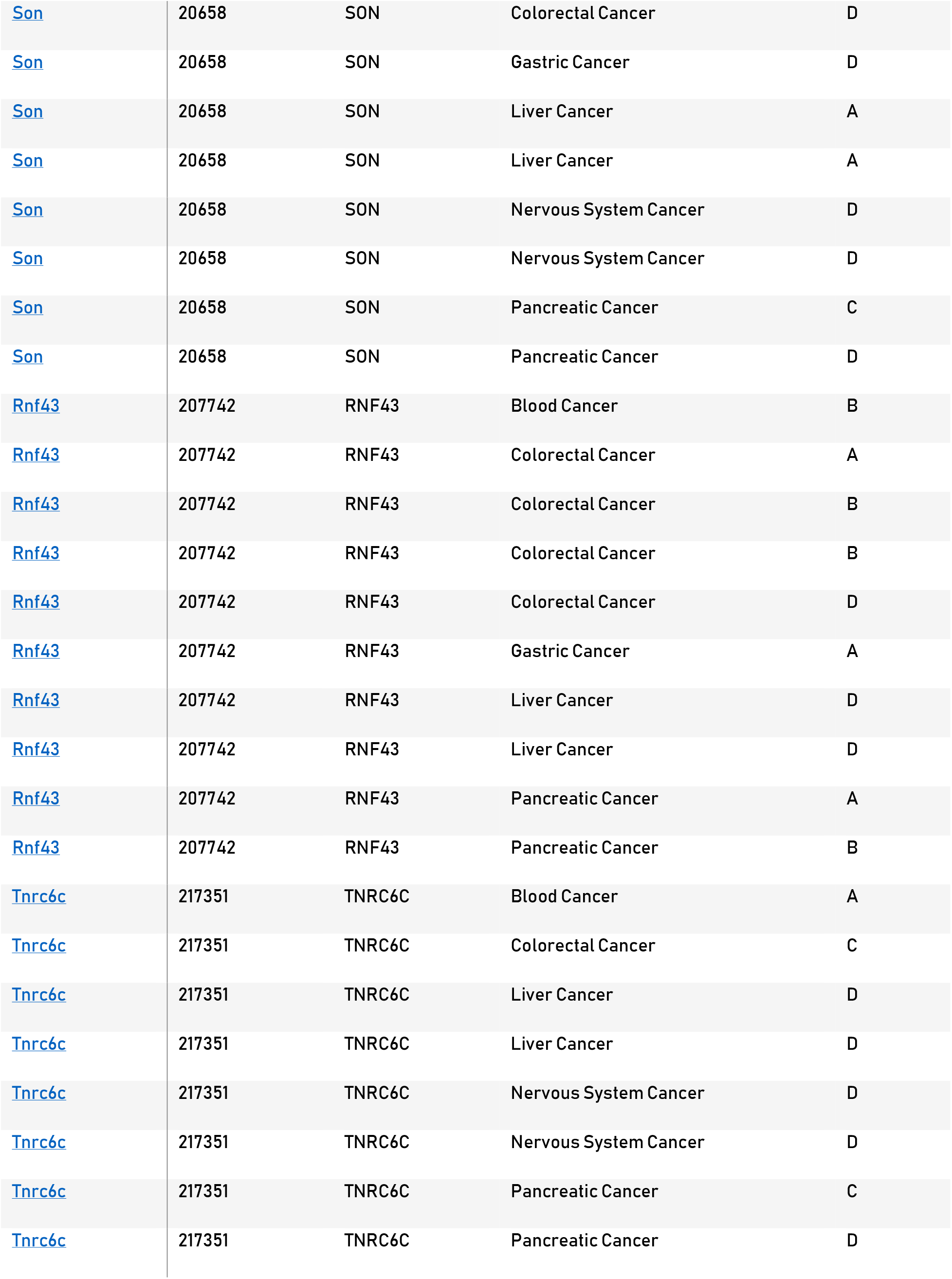
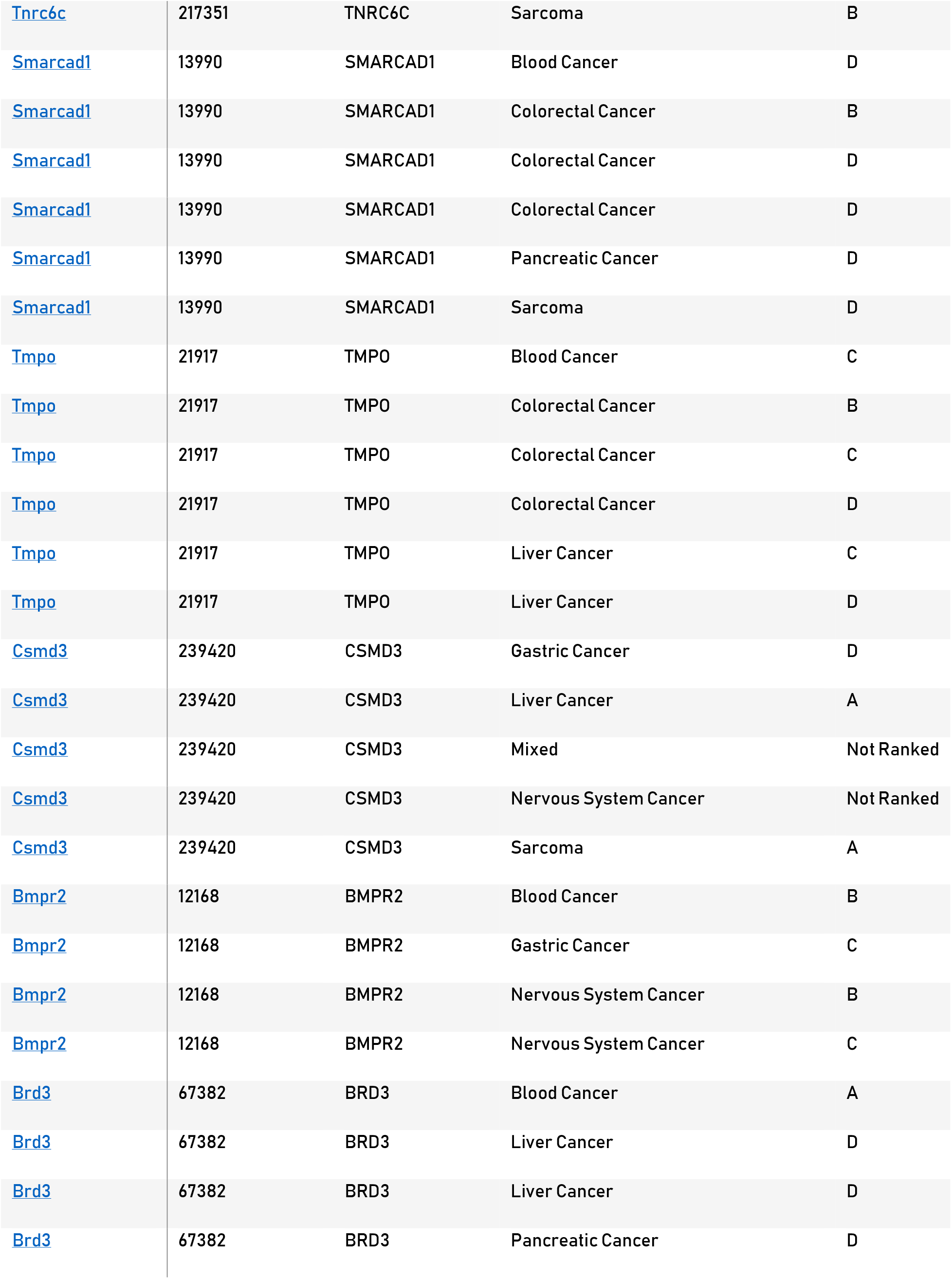
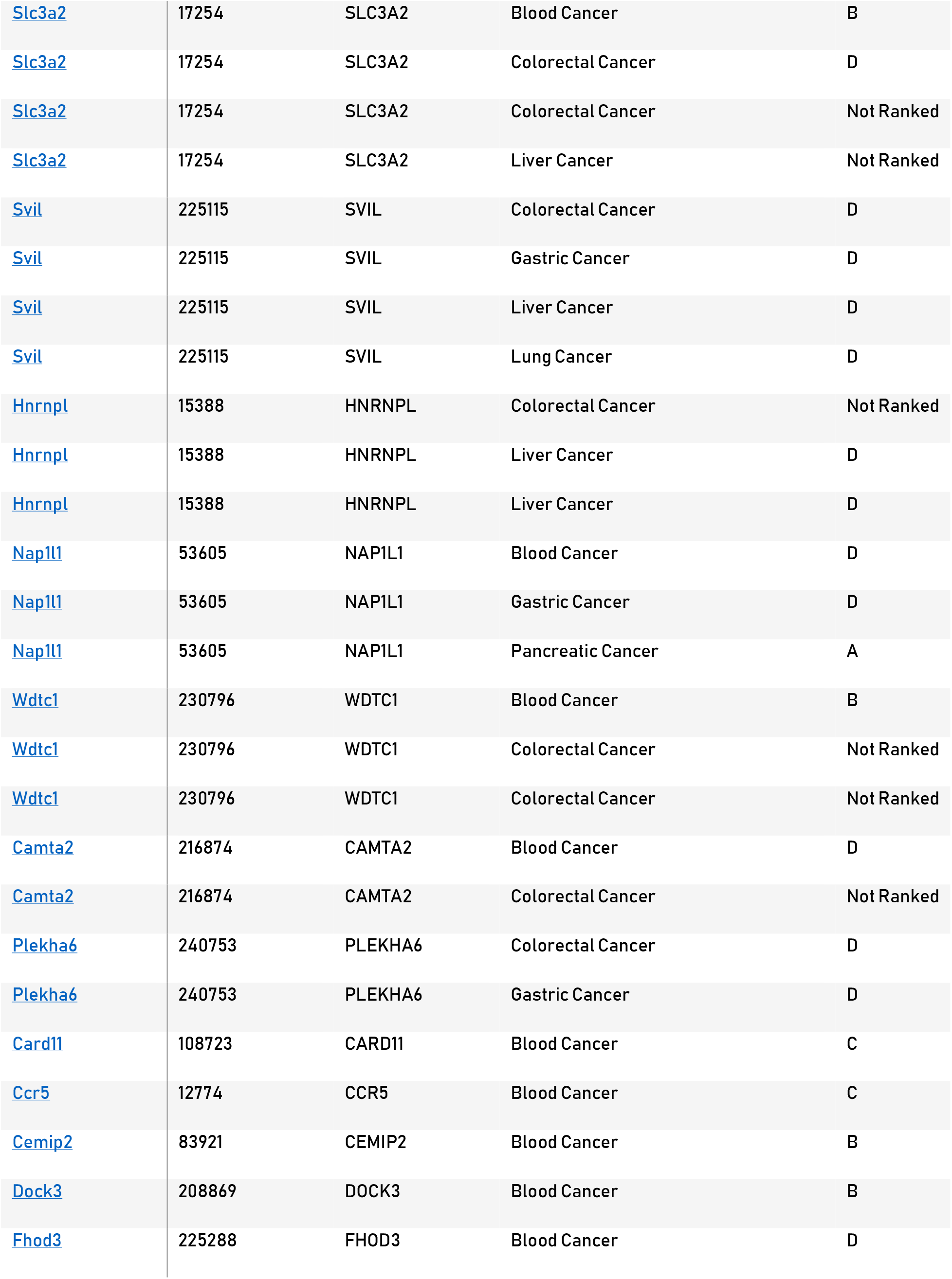
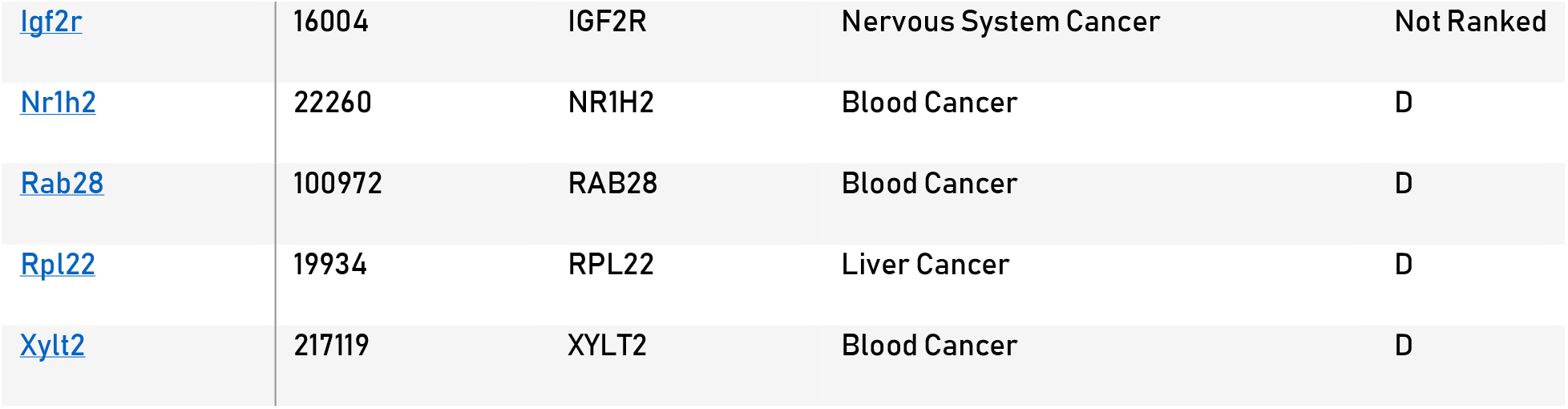
Candidate cancer genes. List with genes described in the candidate cancer gene database that are part of the top 100 most occurring mutations in MSI colon cancer. Rank: Relative rank assigned to CIS in study

## Notes

#### Summary of Updates

Updated author affiliations

## References

1. McGranahan, N. et al. Clonal neoantigens elicit T cell immunoreactivity and sensitivity to immune checkpoint blockade. Science 351, 1463–1469 (2016).

2. Maletzki, C., Schmidt, F., Dirks, W. G., Schmitt, M. & Linnebacher, M. Frameshift-derived neoantigens constitute immunotherapeutic targets for patients with microsatellite-instable haematological malignancies: frameshift peptides for treating MSI+ blood cancers. Eur. J. Cancer Oxf. Engl. 1990 49, 2587–2595 (2013).

3. Brown, S. D. et al. Neo-antigens predicted by tumor genome meta-analysis correlate with increased patient survival. Genome Res. 24, 743–750 (2014).

4. Le, D. T. et al. PD-1 Blockade in Tumors with Mismatch-Repair Deficiency. N. Engl. J. Med. 372, 2509–2520 (2015).

5. Turajlic, S. et al. Insertion-and-deletion-derived tumour-specific neoantigens and the immunogenic phenotype: a pan-cancer analysis. Lancet Oncol. 18, 1009–1021 (2017).

6. Rooij, N. van et al. Tumor Exome Analysis Reveals Neoantigen-Specific T-Cell Reactivity in an Ipilimumab-Responsive Melanoma. J. Clin. Oncol. 31, e439–e442 (2013).

7. Gubin, M. M. et al. Checkpoint blockade cancer immunotherapy targets tumour-specific mutant antigens. Nature 515, 577–581 (2014).

8. Litchfield, K. et al. Escape from nonsense mediated decay associates with anti-tumor immunogenicity. bioRxiv 823716 (2019) doi:10.1101/823716.

9. Kervestin, S. & Jacobson, A. NMD: a multifaceted response to premature translational termination. Nat. Rev. Mol. Cell Biol. 13, 700–712 (2012).

10. Lin, E. I. et al. Mutational profiling of colorectal cancers with microsatellite instability. Oncotarget 6, 42334 (2015).

11. Boyer, J. C. et al. Sequence dependent instability of mononucleotide microsatellites in cultured mismatch repair proficient and deficient mammalian cells. Hum. Mol. Genet. 11, 707–713 (2002).

12. Jiricny, J. The multifaceted mismatch-repair system. Nat. Rev. Mol. Cell Biol. 7, 335–346 (2006).

13. Li, G.-M. Mechanisms and functions of DNA mismatch repair. Cell Res. 18, 85–98 (2008).

14. Guinney, J. et al. The consensus molecular subtypes of colorectal cancer. Nat. Med. 21, 1350–1356 (2015).

15. National Institute of Health. The Cancer Genome Atlas. The Cancer Genome Atlas, National Institute of Health, Bethesda, U.S. http://cancergenome.nih.gov/.”.

16. Wu, M. et al. MSI status is associated with distinct clinicopathological features in BRAF mutation colorectal cancer: A systematic review and meta-analysis. Pathol. - Res. Pract. 216, 152791 (2020).

17. Abbott, K. L. et al. The Candidate Cancer Gene Database: a database of cancer driver genes from forward genetic screens in mice. Nucleic Acids Res. 43, D844–D848 (2015).

18. Lindeboom, R. G. H., Supek, F. & Lehner, B. The rules and impact of nonsense-mediated mRNA decay in human cancers. Nat. Genet. 48, 1112–1118 (2016).

19. Sahin, U. et al. Personalized RNA mutanome vaccines mobilize poly-specific therapeutic immunity against cancer. Nature 547, 222–226 (2017).

20. Jasperson, K. W., Tuohy, T. M., Neklason, D. W. & Burt, R. W. Hereditary and Familial Colon Cancer. Gastroenterology 138, 2044–2058 (2010).

21. Bonsack, M. et al. Performance evaluation of MHC class-I binding prediction tools based on an experimentally validated MHC-peptide binding dataset. Cancer Immunol. Res. (2019) doi:10.1158/2326-6066.CIR-18-0584.

22. Alspach, E. et al. MHC-II neoantigens shape tumour immunity and response to immunotherapy. Nature 574, 696–701 (2019).

23. Kreiter, S. et al. Mutant MHC class II epitopes drive therapeutic immune responses to cancer. Nature 520, 692–696 (2015).

24. Garrido, F., Aptsiauri, N., Doorduijn, E. M., Garcia Lora, A. M. & van Hall, T. The urgent need to recover MHC class I in cancers for effective immunotherapy. Curr. Opin. Immunol. 39, 44–51 (2016).

25. Apcher, S., Daskalogianni, C. & Fåhraeus, R. Pioneer translation products as an alternative source for MHC-I antigenic peptides. Mol. Immunol. 68, 68–71 (2015).

26. Lindeboom, R. G. H., Vermeulen, M., Lehner, B. & Supek, F. The impact of nonsense-mediated mRNA decay on genetic disease, gene editing and cancer immunotherapy. Nat. Genet. 51, 1645–1651 (2019).

27. Pembrolizumab With Ataluren in Patients With Metastatic pMMR and dMMR Colorectal Carcinoma or Metastatic dMMR Endometrial Carcinoma: the ATAPEMBRO Study - Full Text View - ClinicalTrials.gov. https://clinicaltrials.gov/ct2/show/NCT04014530.

28. Cibulskis, K. et al. Sensitive detection of somatic point mutations in impure and heterogeneous cancer samples. Nat. Biotechnol. 31, 213–219 (2013).

29. Koboldt, D. C. et al. VarScan: variant detection in massively parallel sequencing of individual and pooled samples. Bioinforma. Oxf. Engl. 25, 2283–2285 (2009).

30. Shiraishi, Y. et al. An empirical Bayesian framework for somatic mutation detection from cancer genome sequencing data. Nucleic Acids Res. 41, e89–e89 (2013).

31. Li, H. A statistical framework for SNP calling, mutation discovery, association mapping and population genetical parameter estimation from sequencing data. Bioinformatics 27, 2987–2993 (2011).

32. Li, B. & Dewey, C. N. RSEM: accurate transcript quantification from RNA-Seq data with or without a reference genome. BMC Bioinformatics 12, 323 (2011).

33. Zerbino, D. R. et al. Ensembl 2018. Nucleic Acids Res. 46, D754–D761 (2018).

34. McKenna, A. et al. The Genome Analysis Toolkit: A MapReduce framework for analyzing next-generation DNA sequencing data. Genome Res. 20, 1297–1303 (2010).

35. Richards, S. et al. Standards and Guidelines for the Interpretation of Sequence Variants: A Joint Consensus Recommendation of the American College of Medical Genetics and Genomics and the Association for Molecular Pathology. Genet. Med. Off. J. Am. Coll. Med. Genet. 17, 405–424 (2015).

